# Mutation-biased adaptation is consequential even in large bacterial populations

**DOI:** 10.1101/2025.06.16.655099

**Authors:** Jake N. Barber, Alejandro Couce

## Abstract

Adaptive variation is typically presented to selection in a biased manner, owing to vast differences in effective mutation rates across genomes, organisms and environments. Yet, how strongly and how generally such bias shapes adaptation remains poorly understood. Here we used a well-known experimental system in which two *Escherichia coli* mutator variants evolve antibiotic resistance via two mutationally favored, but genetically distinct routes. Through simulations and experiments, we show that the scaling of mutation-biased adaptation with population size is complex, highly dependent on biological details, and—most critically—on how closely mutation bias aligns with selection. Contrary to the common view, we find that mutation-biased adaptation may not wane in large populations but instead intensify depending on the nature of the bias. Crucially, we demonstrate that distinct mutation biases can produce markedly different collateral sensitivity profiles to multiple antibiotics, even at large population sizes. These findings suggest that mutation-biased adaptation may not only be widespread but also have far-reaching and unpredictable consequences both within and beyond the original selective context.

## INTRODUCTION

Understanding the factors that facilitate or constrain the action of natural selection is a fundamental goal in evolutionary biology^1,2^. A central question is how readily available adaptive alleles are across organisms and conditions^3^. Experiments have revealed that the number of loci that can sustain adaptive mutations in a novel environment is typically very large^4–6^. However, this large pool of potentially adaptive variation is rarely presented to natural selection in a uniform manner. This is because beneficial mutation rates vary markedly across organisms, conditions, and even among genes and genetic elements within the same genome^7–10^. Examples include specific local sequence contexts (e.g., short tandem repeats^11^, CpG sites^12^), the mutational specificities of endogenous and exogenous mutagens (e.g., oxidised guanines)^13^, and asymmetries in the genetic architecture of traits that make some elements better suited than others to underpin phenotypic change^5,8^. As a result, the fraction of variation available to selection is often biased toward a few mutationally likely variants. How strongly—and how broadly—this biased supply of heritable variation shapes adaptation remains poorly understood.

The possibility of mutation biases shaping adaptive outcomes (i.e., mutation-biased adaptation) was overlooked by the Modern Synthesis, which emphasized adaptation from standing variation and modelled natural selection as always having the fittest variants at its disposal^14^. However, it is now clear that in many relevant scenarios, populations cannot rely on pre-existing variation for adaptation. A prime example is microbial pathogens, which typically adapt to new hosts and antimicrobial therapies through newly generated variation—either by mutation^15^ or horizontal gene transfer^16^. Under these circumstances, it seems reasonable that the direction and pace of adaptation may be influenced by which mutations arise first.

A first theoretical exploration of this possibility came from Yampolsky & Stoltzfus (2001)^17^, who considered a simple model involving two incompatible adaptive mutations—one common but less fit, the other rarer yet fitter. Across a broad range of parameters, they found that the less-fit mutation often prevails simply because it arises much earlier, precluding the fixation of the superior variant—a phenomenon more pronounced in small populations with stark differences in mutation rates. However, skeptics have criticized the idea of mutation-biased adaptation as limited to exceptional cases, contending that it relies on special conditions—such as strong incompatibilities among beneficial mutations, or population sizes and mutation rates so low that competing mutations never coexist before fixation^18^.

Recent theoretical work has revealed that the restrictive assumptions of early models can be relaxed and mutation-biased adaptation still remain influential^19^. Moreover, mounting empirical evidence now shows that, across multiple model systems, the spectrum of adaptive substitutions consistently mirrors the underlying mutation biases known for each organism and context^20,21^. Examples include laboratory evolution experiments with viruses^20^, bacteria^22^ and yeast^23^, as well as natural cases of parallel adaptation in bacteria^24^, animals^25^ and cancer cells^26^. Overall, these observations suggest a view in which adaptive evolution proceeds not merely by the survival of the fittest, but rather of the likeliest among the fittest.

Still, several open questions remain about the generality and significance of mutation-biased adaptation. One important issue is how influential it is across different population-genetic regimes. In particular, it has long been held that mutation bias should matter less in large populations with high mutation rates^18^. This intuition is readily apparent in the simple two-allele case, where a mutationally-favored but inferior allele prevails only if the superior allele arrives too late^17^. When mutation supply is abundant, the superior allele is more likely to arise in time and outcompete the favored one. The expectation, therefore, scales proportionally: several-fold biases in mutation should translate into several-fold biases in the adaptive outcomes^17^.

The situation becomes more complicated when considering more realistic, multiple-allele models. Sequencing shows that adapting microbial populations often consist of many clones carrying distinct combinations of beneficial alleles that arise and compete in complex ways before any one genotype fixes (i.e., clonal interference)^27–29^. Here, what matters for mutation-biased adaptation is no longer how early a mutationally-favored allele arrives, but rather how early a mutationally-favored *combination* arrives. While simulations confirm that mutation-biased adaptation still weakens with increasing mutation supply in this scenario^19^, the extent of this effect likely depends on idiosyncratic details such as the number and effect sizes of adaptive mutations and the genetic interactions among them (i.e., epistasis)—complications yet to be explored. As a result, how mutation bias scales with mutation supply becomes an empirical question, and comprehensive experimental tests are still lacking.

A second key issue concerns the real-world impact of mutation-biased adaptation. Skeptics have argued that the enrichment of bias-favored alleles observed across multiple model systems may simply reflect a lack of functional effects at these sites, as predicted by Neutral Theory^18,30^—in other words, that we are observing mutation-biased neutral evolution rather than mutation-biased adaptation. While many sites are expected to be neutral and thus to conform to prevailing biases over time, experimental evolution studies clearly demonstrate that many fixed, mutationally-favoured alleles are indeed adaptive^5,20,31,32^. Still, this mutation-driven genetic divergence often results in broadly comparable fitness levels in the original selective environment, suggesting that these divergent genotypes may represent functionally redundant solutions to the same adaptive challenge. Does this imply that mutation bias merely shapes patterns of molecular parallelism, without broader evolutionary consequences?

Two phenomena, recurrently seen in experimental evolution, suggest otherwise. First, genetic incompatibilities occur between mutations, sometimes so strong that the effect of a given mutation can shift across deleterious, neutral or beneficial depending on the background (i.e., sign epistasis)^33^. This context-dependency means that seemingly equivalent initial adaptive steps can quickly channel lineages into divergent trajectories^34^, differing in both the speed and ultimate height of their fitness gains ^35^— potentially even enabling access to initially inaccessible solutions. Second, adaptive mutations often affect multiple traits simultaneously (i.e., pleiotropy)—especially among early adaptations, which frequently target global transcriptional regulators or cause major structural changes in enzymes^36–38^. As a result, populations reaching similar fitness in the same environment can, due to differing pleiotropic profiles, harbor hidden phenotypic variation that may preadapt (or maladapt) them to future challenges^35,39^. Taken together, because mutation bias can either amplify or dampen the effects of epistasis and pleiotropy, mutationally-favored paths are likely to have unpredictable evolutionary prospects—both within and beyond the original selective context.

How readily and how strongly mutation-driven divergence can alter later evolutionary prospects is unknown. These questions bear relevance to many basic and applied problems in biology, such as understanding patterns of biodiversity and speciation^40,41^, and the capacity of populations to cope with new challenges—critical in conservation and in pest and pathogen control^42^. For instance, in the fight against antibiotic resistance, a promising strategy proposes the use of drug combinations or cycling regimens that exploit collateral sensitivity^43,44^—the observation that resistance to one drug sometimes confer hypersensitivity to another. The success of this strategy depends on understanding the factors that determine evolutionary reproducibility^45,46^, as well as on our ability to produce precise predictive models—endeavors that may benefit from accounting for the effects of mutation bias.

To address these questions, we turned to a well-known model system that provides a striking example of mutation-driven molecular divergence. In a prior work^22^, we evolved *Escherichia coli* to increasing antibiotic concentrations, including a wild-type strain and two mutator variants–DNA-repair mutants with strongly biased elevations in spontaneous mutation rate^13^. The distinct mutation biases of the mutators drove the populations along markedly different genetic trajectories, with one mutator mostly exploiting adaptive changes in the antibiotic’s cellular target, and the other favouring changes in a plasmid-borne, antibiotic-degrading enzyme. Here, we first parameterized a simple population-genetics model to reproduce these findings and explore how other factors affect the outcomes. We then repeated the evolution experiment across a 100-fold range of population sizes and found a distinct scaling behaviour of each mutation bias with mutation supply, consistent with our model predictions. Moreover, despite a diminishing influence, mutation bias remained highly consequential, producing different collateral sensitivity profiles even at the largest population sizes.

## RESULTS

### Mutation-biased adaptation shows rich scaling with population size

To explore the sensitivity of mutation-biased adaptation to a range of relevant population genetics parameters, we built a simple simulation model based upon a previously characterised experimental system^22^, so that the model’s most salient predictions can later be tested in the laboratory. The experimental system can be conceived as a two-peak adaptive landscape, where the peaks are similar in height but differ in accessibility (Figure 1A). One peak is reached via a “fast” route, involving mutations in a plasmid-borne, antibiotic-degrading enzyme (TEM-1 β-lactamase); while the other peak is reached via a “slow” route, involving mutations in the cellular target of the antibiotic (PBP3 transpeptidase).

**Figure 1.**
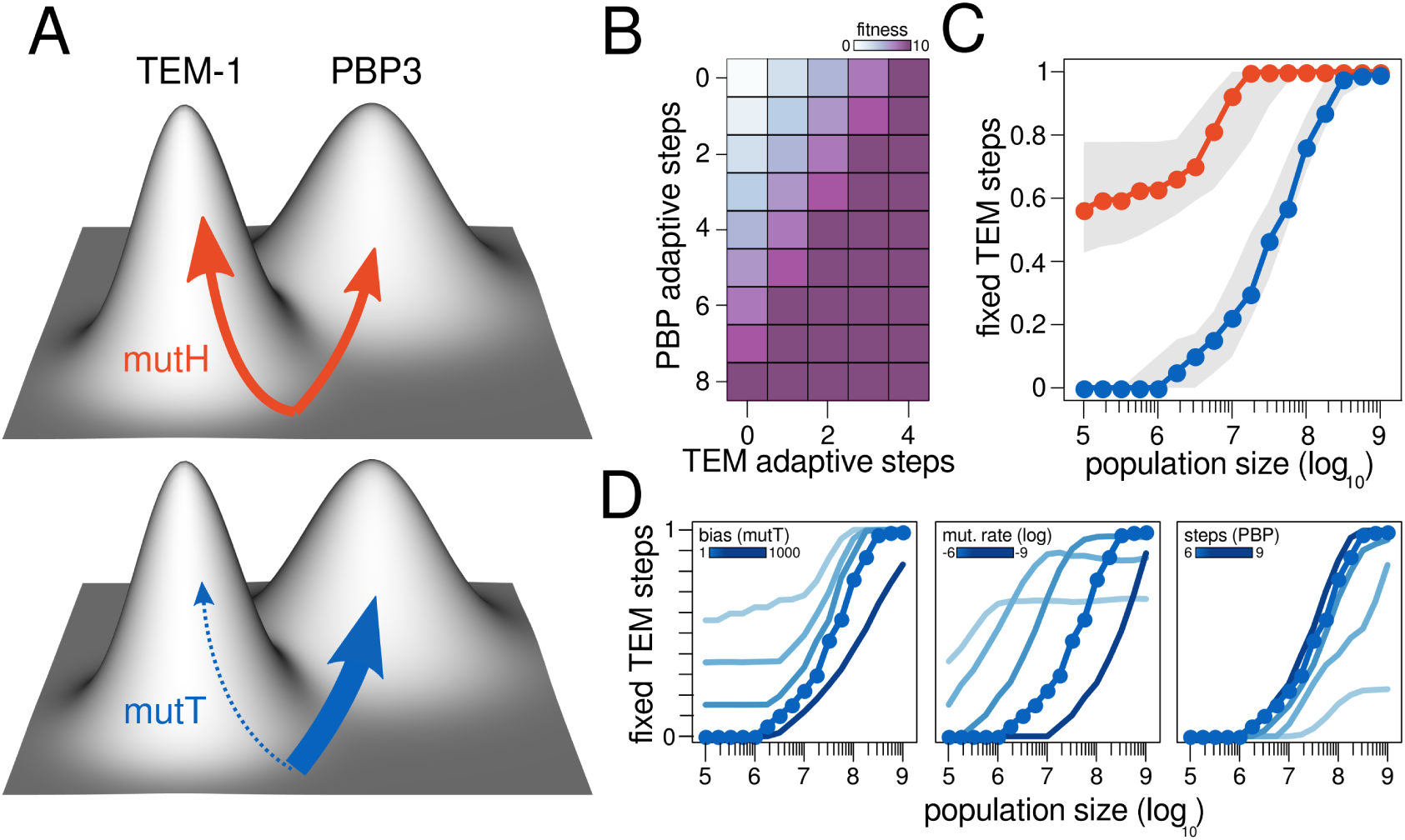
Simulation framework and main outcomes. We built a simple computation model capturing the basic features of the experimental system. (A) We considered an adaptive landscape with two peaks of similar height (fitness) but differing accessibility: one reached via mutations in the plasmid-borne TEM-1 β-lactamase, the other via mutations in the chromosomal transpeptidase PBP3. We simulated two mutators with contrasting spectra: a broad-spectrum mutator (MutH) favoring both paths, and a narrow-spectrum mutator (MutT) biased toward the less accessible PBP3 route. (B) The actual fitness map assumes additive mutation effects up to a fitness ceiling.Since the TEM-1 path requires fewer steps, each mutation yields a larger fitness gain, making this route selectively favored in the absence of mutation bias. (C) Scaling of mutation-biased adaptation with population size. Lines show the median percentage of TEM-1 mutations in high-fitness endpoint genotypes; shaded areas denote interquartile ranges. (D) Sensitivity of the MutT mutator scaling to key model parameters. For reference, the original results from Fig. 1C are highlighted with overlaid dots. Left: shortening the PBP3 path relative to the original setting favors PBP3-driven adaptation; extending it has little effect. Center: weaker mutation bias increase TEM-1 route dominance, stronger ones have little effect. Right: higher baseline mutation rates weaken bias effects, but reduce the dependence on population size.

We sought to keep the model as simple as possible while anchoring core assumptions in relevant biological details. For instance, we assumed fitness peaks of equal height because experimental populations following either route ultimately reached comparable resistance levels^22^. Likewise, we assumed that the TEM-1 route climbs faster to an adaptive peak based on two observations. First, reconstruction experiments show that the most beneficial TEM-1 allele (G238S) confers a resistance increase several-fold higher than the most prevalent, presumably best PBP3 allele (I532L)^47,48^. Second, populations following the PBP3 route required more mutations to reach comparable resistance levels: they contained more changes in the target gene (*ftsI*), as well as more alterations in efflux pumps and permeability^22^.

Another relevant assumption pertains to epistasis. High-resistance genotypes in the experiment often combined varying numbers of PBP3 and TEM-1 mutations^22^. This suggests that, while mild forms of epistasis are likely present, strong genetic incompatibilities are rare among these loci. For simplicity, we therefore assumed that the effects of PBP3 and TEM-1 mutations are purely additive up to a defined fitness maximum (Figure 1B).

We considered asexual populations that, due to distinct mutation biases, explore the fitness landscape either broadly or narrowly. In the original study with *E. coli*, the broad-exploring type corresponded to a MutH-defective mutator, whereas the narrow-exploring one corresponded to a MutT-defective mutator. Both mutators were capable of producing adaptive mutations in PBP3, although each favouring a specific subset of alleles. These subsets likely have different distributions of fitness effects, but we lack information on how they differ and therefore did not consider this possibility. The crucial difference, however, lies in each mutators’s ability to evolve TEM-1. MutH mutators elevate GC→AT and AT→GC transitions, which underlie some of the most beneficial substitutions known in TEM-1. In contrast, MutT mutators elevate AT→CG transversions, which do not to produce any relevant beneficial TEM-1 allele^47^. As a result, the basis for the diverging exploration preferences lies in a coincidental match between each mutator’s mutation biases and the specific nucleotide changes that are beneficial in each target gene.

Simulations confirm the intuition that the influence of mutation bias diminishes with increasing mutation supply (Figure 1C). However, the relationship is not simply linear, instead following a log-sigmoidal shape that depend strongly on the model parameters. For instance, the scaling markedly differs between a broad- and a narrow-exploring mutator (Figure 1C). At large population sizes, both types evolve via the faster, TEM-1 route. But differences become quickly apparent at smaller sizes. The broad-exploring mutator levels off at intermediate values, indicating that populations evolve via mixed trajectories combining TEM-1 and PBP3 mutations. In contrast, the narrow-exploring mutator displays a steeper, all-or-nothing response: its curve spans the full range of outcomes, indicating that populations can sharply shift from adapting exclusively via PBP3 to exclusively via TEM-1.

Variation to key model parameters result into a wide diversity of scaling behaviors (Fig. 1D). Most relevant is the case in which we alter the strength of the narrow-exploring mutator—effectively, the mutational bias favoring the slow versus the fast route (Fig. 1D, left). Weaker biases produce curves that no longer indicate exclusive adaptation via PBP3 at low population sizes; instead, these plateau into mixed trajectories that combine TEM-1 and PBP3 mutations in varying proportions. In contrast, stronger biases simply delay the population size threshold at which adaptation shifts exclusively to the TEM-1 route. Overall, outcomes are more sensitive to reductions than to increases in bias strength. This is relevant because most environmentally-induced mutagenesis likely fall in this range^49–51^, so minor fluctuations in their strength may potentially yield divergent predictions even under similar conditions.

Another relevant parameter substantially altering the scaling behavior is the baseline beneficial mutation rate. At high mutation rates, mutation-biased adaptation weakens— but becomes less sensitive to population size (Fig. 1D, middle). In this regime, the fast TEM-1 route never fully dominates, even in the largest populations, while the slower PBP3 route ceases to be dominant at progressively smaller sizes. Conversely, at low baseline rates, mutation-biased adaptation becomes more pronounced, as well as more sensitive to population size. In other words, these results reveal a trade-off between the prevalence and strength of mutation-biased adaptation as baseline mutation rates vary. Specifically, mutation-biased adaptation is expected to be widespread but weaker when adaptation involves loss-of-function mutations in multiple genes (e.g., disrupting costly catabolic operons), while less common but stronger when adaptation depends on just a few specific sites (e.g., reversion of an auxotrophy).

Changes to other model details show mixed influence on the scaling behavior. One particularly strong influence comes from relaxing the assumption of unequal route lengths: shortening the PBP3 path relative to TEM-1 blurs the distinction between ‘fast’ and ‘slow,’ aligning mutation bias with selection, and favoring PBP3-driven adaptation across all population sizes (Fig. 1D, right; Fig. S1). Relaxing the equal-height assumption reveals a more surprising effect. Elevating the peak of the TEM-1 route shows little influence in the overall behaviour (Fig. S1). In contrast, similar elevations in the PBP3 peak drastically shift outcomes, even eliminating adaptation via TEM-1 entirely (Fig. S1). Finally, changing the absolute magnitude of selection has only a modest effect on outcomes (Fig. S1). Taken together, these results reveal a clear asymmetry: increasing selection for the (already selection-favored) TEM-1 route changes little, but increasing selection for the mutation-favored PBP3 path quickly renders it the predominant adaptive outcome.

### The MutT mutator bias increases extinction risk across population sizes

To experimentally test how mutation supply shapes mutation-biased adaptation, we repeated the original evolution experiment with the two mutators across a 100-fold range of population sizes. We established five regimes—Large (L), Medium-Large (ML), Medium (M), Medium-Small (MS), and Small (S)—using three types of microtiter plates filled to different volumes (Fig. 2A). All other conditions followed the original study^22^ (see Methods), most critically the 1:50 daily dilution, which yields 5.64 generations per passage. The volumes differed by factors of √10 across regimes. This ensures that bottleneck population sizes are evenly spaced across a 100-fold gradient, which we estimated ranged from ∼6.7 × 10⁷ cells (L) to ∼6.4 × 10⁵ cells (S) in the absence of antibiotic. Each mutator population underwent 28 days of daily passages, and antibiotic concentrations were doubled every two passages. At the end of the experiment, cefotaxime concentrations were ∼1,000-fold larger than the minimal inhibitory concentration for the ancestral strains (0.064 mg/L).

**Figure 2.**
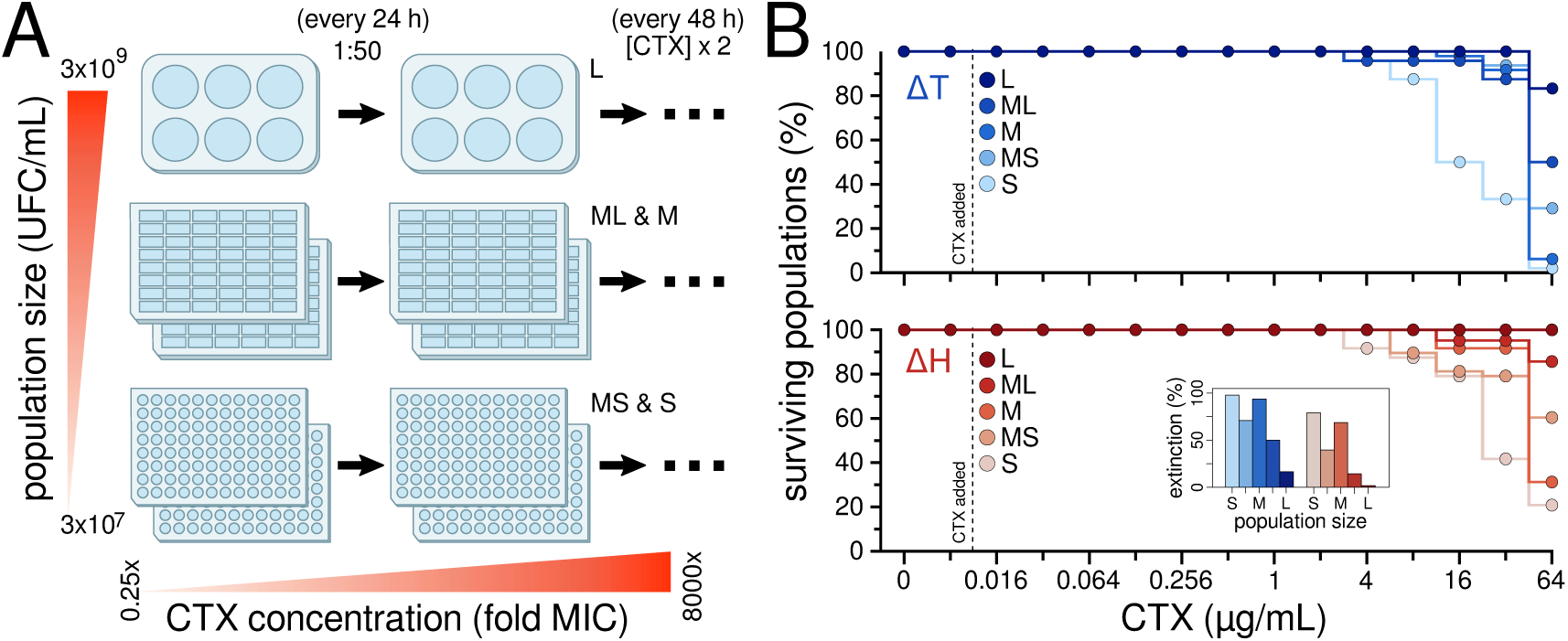
Experimental framework and main outcomes. (A) To explore mutation-biased adaptation across population sizes, we used three different microplate formats to set up five culture volumes covering a 100-fold difference in population sizes in an even manner: Large (L), Medium-Large (ML), Medium (M), Medium-Small (MS), and Small (S). The remaining details were kept as close as possible to the original 2015 experiment. (B) Experiments involved 28 days of 1:50 serial passaging under increasing concentrations of antibiotic, reaching a final concentration ∼1,000-fold larger than the minimal inhibitory concentration for the ancestors. The plots show the percentage of surviving populations across regimes for the two mutators. Inset: in every regime, MutH populations out-survived MutT populations, most likely explained by its preference for the rapid, and therefore safer TEM-1 route.

We found that extinction probability depended strongly on population size: across both mutators, 92% of Large (L) populations persisted until the end of the experiment, compared to just 6% of Small (S) populations (Fig. 2B). This suggests that mutation supply becomes severely limited in the smaller population sizes, with populations more often failing to keep pace with the selective pressure. Most crucially, we observed a systematic survival advantage for MutH-defective mutators over MutT-defective ones (Fig. 2B, inset). In every size class, MutH populations out-survived MutT populations— a pattern unlikely just by chance (P = 0.03, one-sided binomial test). This suggests that in a scenario of steadily increasing lethal selection, the faster route (TEM-1) is also the safest route of adaptation. While a slight MutH advantage was observed in the original study^22^, the broader range of conditions used here reveals the trend more clearly: across population sizes, MutH mutators fare better because they evolve preferentially via the safer adaptive path.

### Mutation biases shapes the genetic basis of adaptation across population sizes

The molecular mechanisms underlying high-level resistance to cefotaxime are well known. In our system, they include two primary routes: (i) point mutations that reduce the drug’s affinity for its cellular target, the PBP3 transpeptidase^52^, and (ii) substitutions that broaden the hydrolytic spectrum of the plasmid-borne β-lactamase TEM-1^47^. Changes in other loci—reducing outer membrane permeability^53^ or increasing efflux^54^— also contribute, particularly at lower drug concentrations^22^. We first aimed to obtain a high-resolution view of how the relative frequency of mutations in TEM-1 versus PBP3 varied with mutation bias and population size, for which we used targeted Sanger sequencing.

We assembled a collection of 75 independent, endpoint isolates: 50 from MutT populations (10 per size class) and 25 from MutH populations (5 per class). We sequenced the loci encoding for TEM-1 and PBP3 (*bla*_TEM-1_ and *ftsI*, respectively) in all isolates. In total, we identified 56 nonsynonymous mutations in *bla*_TEM-1_ (5 unique) and 91 in *ftsI* (17 unique) (Fig. 3A). The most common alleles were *bla*_TEM-1_ G238S (present in 62.5% of *bla*_TEM-1_ mutations) and *ftsI* I532L (found in 38.5% of *ftsI* mutations), both previously known to underpin high-level cefotaxime resistance^47,48^. Of note, only three mutations—A498T in *ftsI* and E104K and G238S in *bla*_TEM-1_—were shared between mutator backgrounds, further evidencing the role of mutation bias in shaping adaptive outcomes.

**Figure 3.**
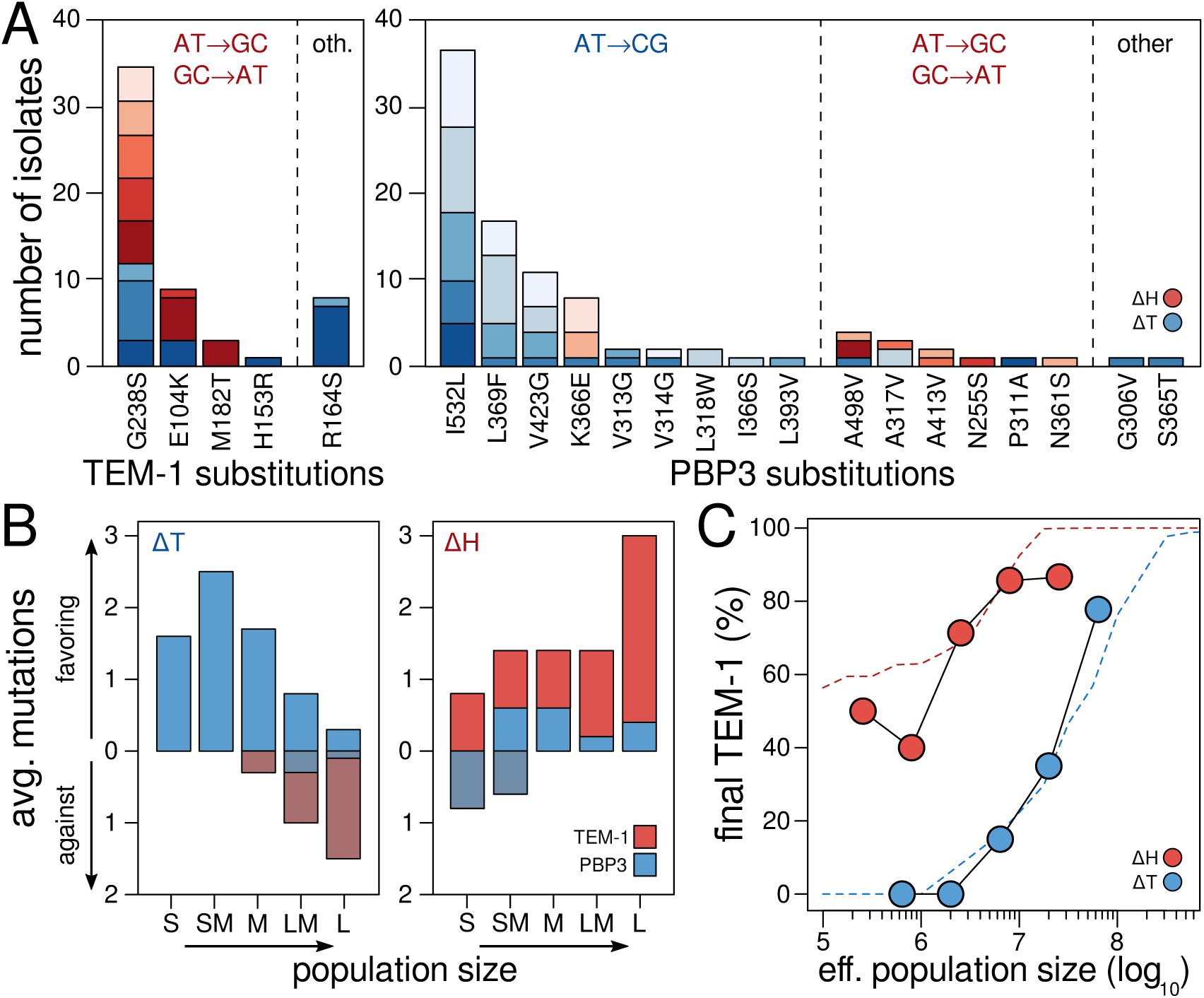
Mutationally-favored substitutions in the focal loci across population sizes. (A) Abundance of different amino acid substitutions among 75 endpoint isolates (50 MutT, 25 MutH). Bars show counts per position in TEM-1 (left panel) and PBP3 (right), grouped by whether the underlying nucleotide change matches each mutator’s bias (indicated on top). Darker colors correspond with larger population sizes. (B) Average number of bias-matching substitutions per isolate for each mutator (MutT, left; MutH, right) across population-size classes. In MutT, mutation-bias adaptation declines with increasing size: small populations fix only bias-favoured PBP3 substitutions, whereas non-favoured TEM-1 mutations appear at larger sizes. By contrast, bias-matching substitutions rise with size in MutH, indicating a synergistic effect when the mutator’s bias aligns with selection for optimal mutations in both genes. (C) Empirical scaling of mutation-biased adaptation with population size. Lines show the percentage of TEM-1 mutations in high-fitness endpoint genotypes. Dashed curves represent simple, best-fit simulations. (see text for details).

In line with the original experiment^22^, the two mutators displayed contrasting preferences in their mutational paths: MutH mutators favored *bla*_TEM-1_ over *ftsI* mutations (34 vs. 16), whereas MutT mutators favoured *ftsI* over *bla*_TEM-1_ mutations (24 vs. 75). This trend reflects the predominance of GC→AT and AT→GC transitions among beneficial mutations in *bla*_TEM-1_ (favored by MutH), and AT→CG transversions in *ftsI* (favored by MutT). Notably, the preference for one gene over the other varied with population size in the MutT group, but not so much in MutH (Fig. 3B). In MutT populations, *bla*_TEM-1_ mutations were recovered only in the intermediate to large size regimes, and entirely absent in the smaller population size groups. In contrast, MutH populations consistently harbored mutations in both *bla*_TEM-1_ and *ftsI*, regardless of population size (Fig. 3B).

We next asked how closely the spectrum of adaptive substitutions matches each mutator’s bias across the five population sizes. For MutT lineages, we expected the match to decline steadily with size, because larger populations favor the TEM-1 route, and the MutT bias does not match any relevant mutation along that route^47^. However, the relationship proved more nuanced (Fig. 3B). The proportion of bias-matching substitutions peaked in the medium-small class, and even in the largest populations, a small but significant fraction of bias-matching PBP3 mutations persisted alongside TEM-1 mutations. In contrast, in MutH lineages, the bias-matching fraction of mutations increased with population size, reaching 100% from the intermediate class upward. Moreover, we observed a sharp rise in the number of bias-matching mutations in the Large (L) size regime. This likely reflects that the MutH bias produces several substitutions that when combined confer high beta-lactam resistance (i.e., G238S, E104K and M182T, the combination forms the clinically important TEM-52)^55^; suggesting synergistic effects when a mutator’s bias aligns with the selection-favored pathway.

Finally, we examined whether the empirical trends align with the simulation’s predictions regarding the prevalence of TEM-1 and PBP3 mutations across population sizes and mutator backgrounds. A key challenge in this comparison is estimating the effective population size (Nₑ) for the experiments. In a serial passaging regime, Nₑ can be approximated as the product of the bottleneck size (N₀) and the number of generations per passage (g)^56^, such that Nₑ = N₀ × g. In our case, this yields a range of ∼3.6 × 10⁶ to ∼3.8 × 10⁸ cells across our Small to Large regimes. However, these values represent an upper bound, as they do not account for the recurrent population declines that occur each time the antibiotic concentration doubles. We therefore opted to adjust N₀ to obtain values within the range used in the simulations. After this adjustment, the shape of the MutT curve showed a remarkable match with that obtained from the simulation model (Fig. 3C). For the MutH mutator, the adjustment required a further modification in the model, namely a relaxation of the assumption that it favors both routes equally: simply making that its bias slightly favors the PBP3 route over the TEM-1 pathway sufficed.

### MutT mutators require more second site mutations to keep up with adaptation

To assess whether the patterns of mutation-biased adaptation observed in our focal genes extended to secondary loci genome-wide, we selected five independent isolates per mutator background and population size class (10 groups, 50 isolates total) and subjected them to whole-genome sequencing. Given the large number of mutations expected in mutator strains, we focused our analysis on genes hit in more than four groups and those for which prior literature supports a link to β-lactam resistance (Methods).

The results show a great diversity of mutations in genes encoding efflux pumps, porins, and associated regulatory elements. This is consistent with previous studies showing that β-lactam resistance often involves a combination of increased enzymatic degradation, reduced PBP affinity, enhanced efflux, and decreased permeability^22,27,53^. The locus with most hits across treatments was *acrB* (18 hits, 10 in MutT and 8 in MutH lineages), which encodes the inner membrane component of the AcrAB-TolC efflux pump^54^. Consistent with this, the second most commonly hit was *acrR* (10 hits, all in MutT populations), which encodes the major transcriptional repressor of the same efflux pump. The third locus with most hits is *ompR* (8 hits, 7 in MutT populations), encoding a positive regulator controlling the expression of the major porins through which β-lactam enter the cells, OmpF and OmpC^57^.

While both mutators broadly adapted through the same genetic targets, signatures of mutation-biased adaptation were apparent across secondary loci. First, several key accessory genes were mutated exclusively in MutT populations and never in MutH populations (i.e., *acrR*, *envZ*, *ompF*, and *marR*) (Fig. 4A). Second, at the allele level, there was a striking lack of overlap among backgrounds, with only 6 shared alleles out of 71 cases across MutT and MutH populations (8.5%; *acrB*: 1, *hrpB*: 1, *metK*: 1, *ykfM*: 3; Fig. S2). This mirrors the pattern in *ftsI*, where each mutator accessed a distinct subset of alleles (Fig. 3A). Finally, MutT populations showed both a greater diversity and a higher number of accessory gene mutations than MutH ones (49 vs. 16 distinct alleles, 2.7 vs. 0.73 mutations per genome) (Fig. 4B). This is consistent with the higher extinction risk observed in MutT lineages; supporting the idea that, because they preferentially evolve through the slower—and thus riskier—PBP3 route, MutT mutators can only keep up with strong selection pressures by accumulating more mutations in secondary loci.

**Figure 4.**
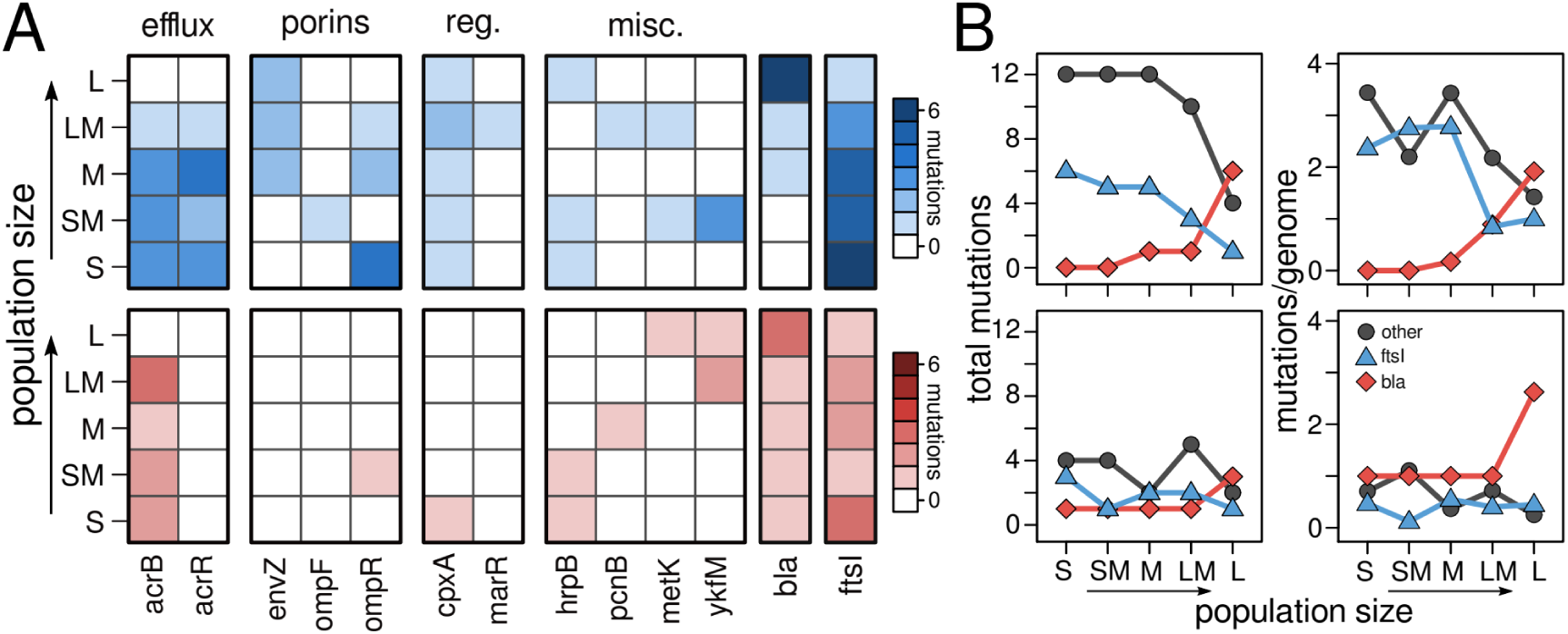
-Signatures of mutation bias across secondary loci. (A) Heatmap showing the distribution of mutations in accessory genes associated with β-lactam resistance across regimes. Several loci (*acrR, envZ, ompF,* and *marR*) are mutated exclusively in MutT populations; only 6 of 71 alleles are shared between mutators (Fig. S2). (B) MutT populations carry both a greater diversity (left) and a higher average number (rigth) of accessory-gene mutations than MutH, consistent with the need to compensate for their reliance on the slower, PBP3-based adaptive route.

### Collateral sensitivities to other antibiotics diverge even at large population sizes

We next asked whether the observed mutation-biased adaptation—evident even at large population sizes—can have real-world consequences. Understanding how mutation bias shapes phenotypes beyond the immediate selective context is key for strategies that rely on trade-offs, such as exploiting collateral sensitivity in pathogenic bacteria^43,44^. Collateral sensitivity typically arises as a by-product of mutations in transcriptional regulators or multidrug resistance systems—alterations we frequently observed in the sequenced isolates (Fig. 5A). While the specific alleles substituted in these elements varied markedly with mutation bias, both mutators broadly adapted through a combination of PBP3 alterations, increased β-lactamase activity, enhanced efflux, and reduced permeability. This broad phenotypic convergence raises the question of whether different mutation biases can still produce distinct collateral sensitivity profiles, particularly at larger population sizes.

**Figure 5.**
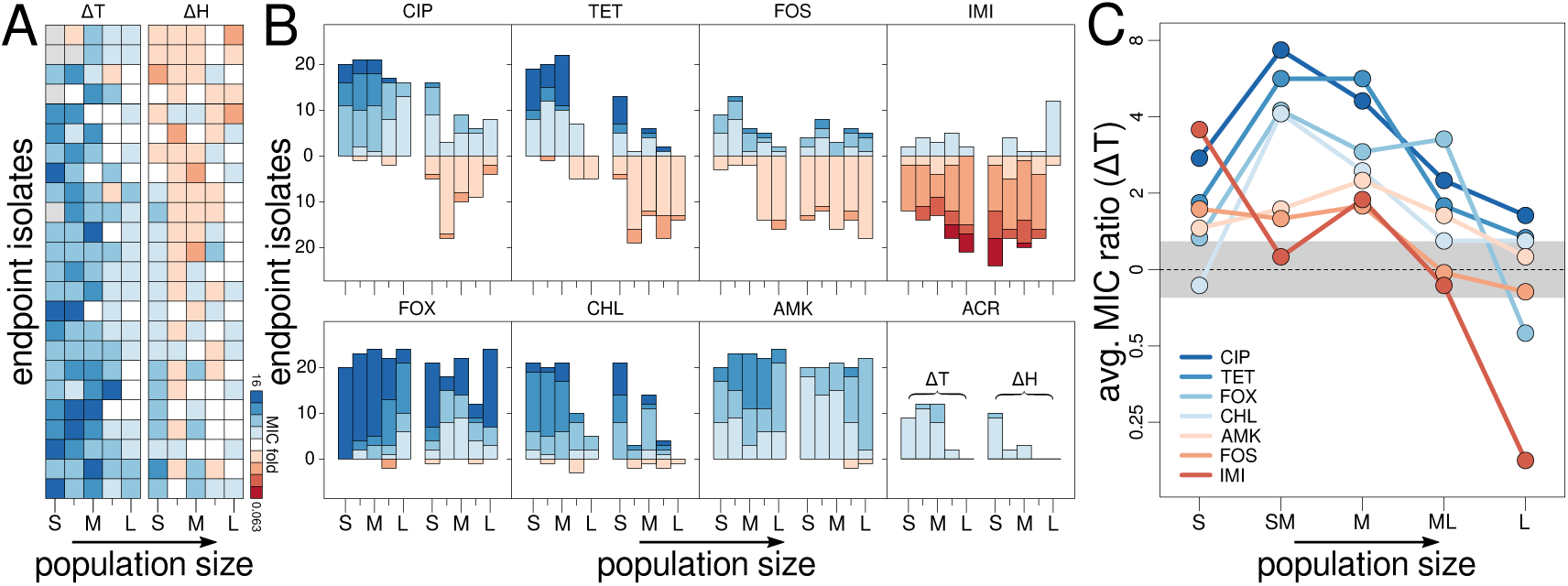
Mutation bias drives collateral-sensitivity divergence across population sizes. (A) Heatmap showing changes in ciprofloxacin susceptibility across population-size classes in 24 endpoint isolates per mutator (MutT, left; MutH, right). Colors indicate fold-changes in MIC relative to the ancestor (white, no change; red, increased susceptibility; blue, increased resistance). **(B)** Summary of MIC fold-changes for all tested antibiotics. Across compounds, MutT isolates consistently evolve higher resistance levels than MutH. **(C)** Values represent the average MIC ratio between mutators, with 1 indicating no difference. Shaded area indicates the 95% confidence interval based on within-treatment variability. Contrary to expectations that bias effects fade with size, divergence peaks in the medium-small (MS) and medium (M) classes. Notably, at the largest sizes, some drugs (e.g., FOX, IMI) show opposite trends, with IMI shifting from resistance in MutH to sensitivity in MutT.

We measured collateral resistance and sensitivity to eight compounds across all treatments, including seven antibiotics from major classes and one antiseptic (Fig. S3). Overall, we found widespread alterations in collateral resistance and sensitivity across all eight drugs, with 71.5% of isolates showing changes in either direction (ranging from 92.9% for amikacin to 22.9% for acriflavin). Crucially, the direction and magnitude of these changes varied markedly by mutator background. Ciprofloxacin and tetracycline provide a particularly illustrative example: most MutH lineages became more sensitive to these drugs (38% and 58%, respectively), whereas most MutT lineages became more resistant (83% and 59%; Fig. 5). Moreover, we noticed that across compounds MutT populations tended to evolve higher resistance than MutH ones (P < 10^-4^, one-sided binomial test)—a pattern evident even when both backgrounds mostly show cross-resistance (e.g., cefoxitin; Fig. 5B). This trend is consistent with the notion that MutT populations require more secondary mutations to compensate for their poor evolvability via the TEM route. Taken together, these findings show that identical selective pressures can yield opposite and distinctive collateral effects depending on the prevailing mutation bias.

Finally, we assessed how mutation-driven differences in collateral sensitivity profiles vary with population size. As mutation bias plays a stronger role in smaller populations, we expected divergence in collateral sensitivity profiles to decline monotonically with size. However, divergence between the mutator profiles instead peaks at the intermediate-small and intermediate size classes (Fig. 5C). Moreover, at the two largest sizes, we observed clear exceptions to the trend of MutT populations becoming more resistant overall (e.g., cefotaxime and imipenem, Fig. 5C). The effect was especially striking for imipenem, where not only the magnitude but also the sign of the effect reversed, with most MutH populations evolving resistance, and most MutT populations evolving susceptibility. These results underscore the limits of collateral sensitivity strategies under strong mutational biases—whether due to mutator strains or environmental factors—as the success of these strategies depends on side-effects remaining robust to common sources of biological and environmental variation^45,46^.

## DISCUSSION

Here we set out to explore how often and how strongly mutation bias shapes adaptation, using a well-established model in which two mutators evolve antibiotic resistance via divergent paths^22^. A first motivation was to test whether mutation-biased adaptation remains relevant at large population sizes, where it is commonly assumed that selection coefficients are the sole determinants of adaptive outcomes^18^. Contrary to this view, our results show that while mutation-biased adaptation may wane in some cases, it can also persist—and even intensify—with population size depending on the nature of the bias.

How can we rationalize these counterintuitive findings? The discussion around mutation-biased adaptation has often focused on the most striking case—when suboptimal yet mutationally-favored variants prevail^17,18^. However, mutation bias can also favor variants that are as fit, or even fitter, than their competitors. In these cases, the net result remains the same: mutationally-favored variants are enriched beyond what selection coefficients alone would predict. The key difference is that, in large populations, mutation-biased adaptation is no longer expected to fade: it either becomes insensitive when equally fit variants are favored, or intensifies when mutation bias aligns with the fittest ones.

Our findings illustrate the range of possible outcomes. Mutation-biased adaptation weakens with size when the bias favors less-fit variants—likely the case for PBP3 mutations, which are less effective for rapid adaptation than TEM-1 mutations. Accordingly, in MutT populations, which favor the PBP3 route, the fraction of fixed mutations matching the bias declined with size (Fig. 3B). By contrast, when the bias favors alleles that are equally or more fit, the pattern reverses: in MutH populations, which favor TEM-1 mutations, bias-matching variants peaked at the largest size (Fig. 3B). Overall, these results suggest that the impact of mutation bias in large populations depends critically on its alignment—or misalignment—with selection coefficients.

Our second motivation was to assess the potential real-world consequences of the mutation-biased divergence we observed across population sizes, by quantifying how the evolved lineages differed in collateral resistance and sensitivity to eight additional antimicrobials. Across compounds, MutT populations generally evolved higher resistance than MutH populations, consistent with their need for additional secondary mutations to compensate for limited access to the TEM-1 route. Most importantly, both the direction and magnitude of collateral effects differed substantially between mutator backgrounds—sometimes so starkly that the profiles were reversed (e.g., ciprofloxacin and tetracycline, where most MutH populations became susceptible, while most MutT ones became resistant) (Fig. 5). Finally, contrary to the expectation that mutation-biased divergence should decrease with population size, divergence peaked at intermediate sizes, with notable differences persisting even at the largest size class. Together, these findings show that mutation bias can steer populations toward divergent collateral-sensitivity profiles, even when evolving from nearly identical backgrounds, with similar population sizes, and under the same selective conditions. This suggests that accounting for mutation bias is key to ensure that strategies that exploit collateral sensitivity in microbial infections achieve success across clinical conditions.

Our study provides a clear example of how mutation bias can shape phenotypes across a wide range of population sizes. To what extent do these findings hold in other systems? Our simulation model, which broadly reproduces the experimental data, suggests that consequential mutation-biased adaptation may be common. Mutators are a relevant experimental model as they readily evolve in asexual populations under moderate to strong selection^58^, and genomic evidence suggests they rise and fall recurrently in natural lineages^59^. Mutators, however, mark only one end of a wider range of scenarios. Many lifestyle-associated stresses can create strong mutation biases in a dose–response manner^9,51^. Examples include starvation^50^, UV light^60^, oxidative stress^61^, heavy metals^62^, and antibiotics^63^. Crucially, our simulations indicate that mutation-biased adaptation is particularly sensitive to variations in bias strength within the several- to tens-fold range. Because lifestyle-associated mutagenesis typically falls within this range, even modest fluctuations in mutagen exposure could steer populations along divergent evolutionary paths under otherwise similar conditions. Even in closely related organisms, the same locus may differ in architecture or contain distinct mutational hotspots—features that can readily account for several- to tens-fold variation in mutation bias^8,32^.

Finally, our simulations reveal that variation in model assumptions—beyond bias strength alone—can produce a rich variety of outcomes. For instance, we found that the strength and population-size dependency of mutation-biased adaptation often trade-off across conditions. A key factor is the baseline mutation rate, which are known to vary substantially across organisms^10^, environments^64^, and mutational targets^8^. Likewise, whether bias favors more-fit or less-fit adaptive variants strongly shapes how outcomes scale with population size. In nature, such alignment between mutation and selection is expected to occur in all range of combinations. Together, these sensitivities suggest that mutation-biased adaptation will show a rich and complex variety of behaviours across biological systems—even before adding further layers of complexity such as multi-peaked landscapes, recombination, or shifting environmental conditions. We hope our results encourage further exploration of these dynamics in a wide range of experimental and natural settings.

## METHODS

### Bacterial strains and media

All strains are derivatives of the *E. coli* K-12 laboratory strain AB1157, with genetic modifications described in Couce *et al.* (2015)^22^. Briefly, the gene encoding the β-lactamase AmpC, despite being normally silent, was deleted to prevent interference with the experimental system. Deletions of *ampC* and the mutator genes *mutT* and *mutH* were introduced via P1 transduction from the Keio collection, followed by pCP20-mediated excision of the kanamycin resistance cassette^65^. All strains carried the *bla*_TEM-1_ gene on plasmid pBRACI, a pBR322 derivative lacking the tetracycline resistance cassette (removed via AatII and AvaI digestion). Bacteria were grown in M9 minimal medium supplemented with 1% glucose and 1% acid hydrolysate of casein, to support maximal growth in the microwell plate format used in the original study. Cefotaxime was purchased as the commercially available powder for injectable solution (Generic Pharmaceutical Equivalent) from Farmacia Santaolalla (Spain) and stored as a 10 mg/mL stock solution at –20 °C.

### Experimental evolution

Populations were serially propagated under static conditions at 37 °C, doubling the concentration of cefotaxime every 48 hours over 28 days. Cultures were diluted 1:50 into fresh medium every 24 hours, yielding ∼5.64 generations per passage. The first two passages were drug-free. On day three, cefotaxime was introduced at 0.016 mg/L (1/4 × MIC, Minimal Inhibitory Concentration; 0.064 mg/L for the ancestral strain), and doubled every two transfers until reaching 64 mg/L (1024×MIC). To generate a ∼100-fold gradient in population size, we used microtiter plates filled to volumes differing by factors of √10. Five regimes were established: Large (L), Medium-Large (ML), Medium (M), Medium-Small (MS), and Small (S). We propagated 48 parallel populations per treatment, except for the largest regimes, where 24 were used to balance manageability with the lower expected extinction risk^27^. Group L was cultured in custom 3D-printed polypropelene wells (Verbatim), designed using an OpenSCAD and printed using a Prusa MK3S+ ; ML and M in 48-deep-well plates (VWR International); and MS and S in 96-well plates (Fisher Scientific). Glycerol stocks were collected every two days and stored at –80 °C. Population survival was tracked by plating 5 μL onto LB agar every 48 hours and visually scoring for presence/absence of bacterial growth.

### Sanger sequencing

Endpoint isolates were obtained from 10 randomly selected MutT-mutator and 5 MutH-mutator populations per size class, chosen among the lineages that survived the longest in the experiment. To assess mutations in the *bla*_TEM-1_ gene, single colonies were streaked onto LB agar, and plasmid DNA was extracted using the GeneJET Plasmid Miniprep Kit (Fisher Scientific). The *bla*_TEM-1_ gene was amplified by PCR using primers 5′-AAG GAT CTT CAC CTA GAT CC-3′ and 5′-CAT TTC CGT GTC GCC CTT ATT C-3′. To assess mutations in *ftsI*, the chromosomal gene was amplified using primers 5′-CAA ATG CAG CAT GTT GAT CC-3′ and 5′-CTA CAG CTA CAA AGA GAT CGC C-3′. Due to the length of this locus (∼1.8 kb), two internal primers—at ∼600 bp (5′-AAG TGA CTG CTC ACC TCA TCG G-3′) and ∼1200 bp (5′-AGC GTT AGT AGA TAC TTA CTC ACG-3′)—were also used for sequencing. Sequencing was performed by Macrogen (https://dna.macrogen.com), and alignments were carried out using MAFFT v7 (https://mafft.cbrc.jp/alignment/software).

### Whole genome sequencing analyses

For each population size class and mutator background, we selected five endpoint isolates, each originating from a different, randomly chosen population. Single colonies were grown overnight in 5 mL LB broth, and genomic DNA was extracted with the DNeasy UltraClean Microbial Kit (Qiagen, Germany). DNA from the five isolates in each treatment was combined into a single, equally weighted pool for deep sequencing. To this end, individual DNA concentrations were quantified with a Qubit 4 fluorometer (Invitrogen, USA), and volumes were adjusted so that each clone contributed the same mass of DNA to the pooled sample. Sequencing services were provided by Novogene (Germany), and performed using Illumina Novaseq 6000 with 150 bp paired-end reads. Each sample was sequenced to a depth of 300x. Variants were called using Breseq v0.33.2 on the population sample setting on default parameters^66^. To minimize false positives, we retained only single-nucleotide polymorphisms present at ≥10 % frequency—twice the standard 5 % threshold recommended for Breseq’s polymorphic mode. Reads were aligned to the fully annotated *E. coli* AB1157 reference genome (NCBI accession: PRJNA731904). All sequencing data have been deposited in the NCBI Sequence Read Archive (accession TBD).

### Antimicrobial susceptibility testing

At the end of the evolution experiment, 24 populations were sampled from each regime (n = 240 total) for characterization of cross-resistance and cross-sensitivity profiles against eight antimicrobials. A single colony was isolated from each population by streaking onto LB agar plates. Colonies were then grown overnight in liquid medium and stored at –80 °C. To perform the susceptibility assays, glycerol stocks were revived overnight, diluted 1:100, and subcultured for 4 hours. Using a 96-pin replicator, cultures were then spotted onto square LB agar plates containing two-fold serial dilutions of each antimicrobial. Minimum inhibitory concentrations (MICs) were recorded in triplicate as the lowest drug concentration that yielded no visible colonies. Tetracycline, fosfomycin, amikacin, acriflavine, chloramphenicol and imipenem were purchased from Fisher Scientific (Spain). Ciprofloxacin was purchased from Merck (Spain), cefoxitin from MedChemExpress (EU) and fosfomycin from Farmacia Santaolalla (Spain). All compounds were stored as stock solutions at –20 °C. Chloramphenicol was dissolved in ethanol, and all others were dissolved in water.

### Computer simulation

We considered adaptation in an asexual, haploid population facing a novel environment using a Wright–Fisher framework—that is, with a fixed population size (N_e_) and discrete, non-overlapping generations. Genotypes are defined by two loci corresponding to the two main resistance routes observed experimentally: one representing mutations along the TEM-1 pathway and the other along the PBP-3 pathway. Mutations from both loci combine additively up to a defined fitness maximum; epistasis is otherwise ignored. In the default configuration both routes ultimately reach the same maximum fitness, but because the TEM-1 path comprises fewer steps, each TEM-1 mutation confers a larger average fitness increment. Hence, in the absence of mutation bias, selection alone would favour the TEM-1 route. The model is class-based: individuals with identical genotypes are pooled, greatly reducing computation time. Each generation proceeds in two stages: (i) Reproduction: for each genotype, its expected contribution to the next generation is calculated in proportion to its current abundance multiplied by its fitness. The entire population is then resampled to exactly N_e_ individuals using these weighted contributions, so fitter genotypes are more likely to expand. (ii) Mutation: after reproduction, we implement mutation using a Poisson-distributed pseudorandom number generation that takes as arguments the basal mutation rate and the number of individuals per genotype. To model mutator bias, the baseline expectation is multiplied by a bias coefficient that inflates mutations toward the favoured pathway (in the default configuration the coefficient is 1 for the TEM-1 route and 100 for the PBP-3 route in MutT backgrounds). Back-mutation and deleterious mutations are ignored. Simulation stop when at least 50 % of the population reaches the maximum fitness class. For each replicate we recorded the proportion of evolutionary steps that followed the TEM-1 pathway (TEM %). Results were averaged over 100 independent replicates. Basic codes are publicly available at https://github.com/ACouce/TBD.

### Statistical analyses and visualization

Both statistical analyses and simulations were conducted in R (version 3.6.3) using built-in functions. One-sided, exact binomial tests were done using the *binom.test()* function. Data visualization was also performed using base R functions, except for Figure 1A, which used the “rgl” package, and Figures 4A and 5A, which used the “pheatmap” package.

## DATA ACCESSIBILITY

All data necessary to replicate the findings will be made publicly available upon publication, either through public data repositories (TBD) or as part of the electronic supplementary material (TBD).

## AUTHORS’ CONTRIBUTIONS

**JB:** Conceptualization (equal); methodology (lead); investigation (lead); formal analysis (supporting); writing – original draft (supporting); writing – review and editing (equal), funding acquisition (supporting) **AC:** Conceptualization (equal); formal analysis (lead); writing – original draft (lead); writing – review and editing (equal), funding acquisition (lead), supervision (lead).

## ACKNOWLEDGMENTS

We thank Lewis Grozinger for assistance with the custom 3D-printed multi-well plates.

## SUPPLEMENTARY FIGURES

**Figure S1.**
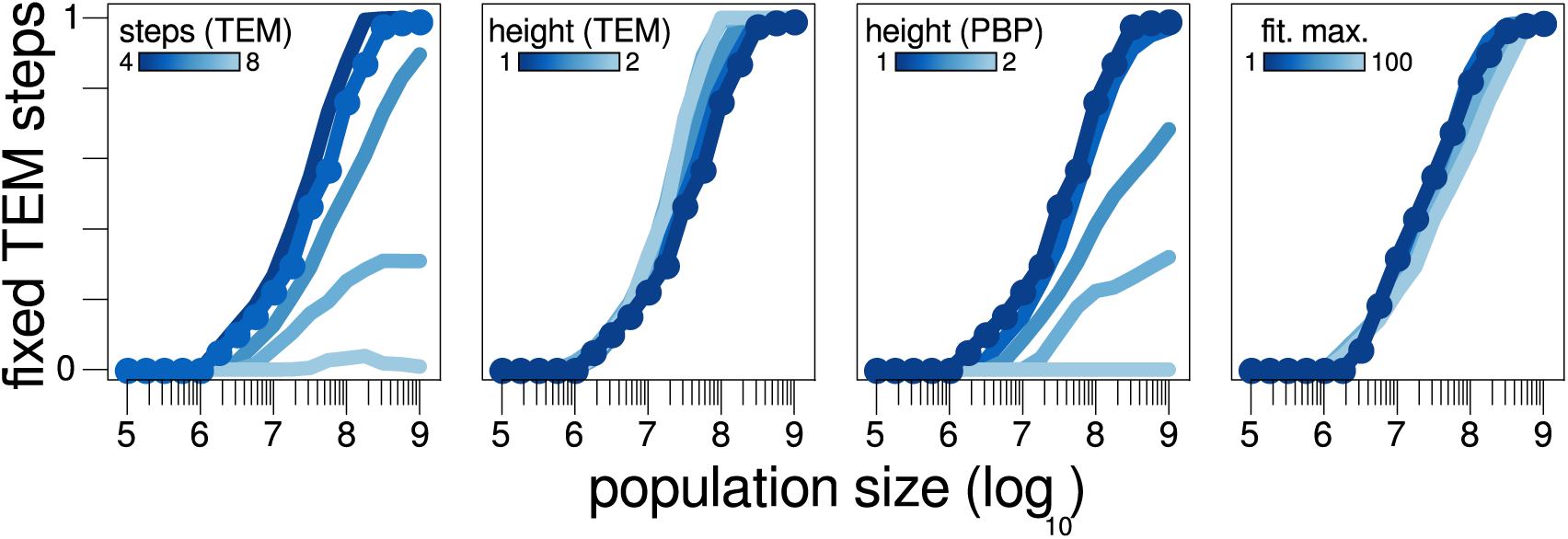
Changes in simulation framework and their effects on adaptive outcomes. Plots show how the probability of mutation-biased adaptation scales with population size under varying model assumptions. The original simulation results for a MutT mutator (main text) are highlighted with overlaid dots. Left: Extending the length of the TEM-1 route relative to the original setting favors PBP3 adaptation; shortening it further has minimal effect. Center-left and center-right: Increasing the height of the TEM-1 peak has little impact, but raising the PBP3 peak strongly shifts outcomes toward the PBP3 route. Right: Increasing the absolute magnitude of selection (per-step fitness gains) has only a modest influence on the adaptive trajectory.

**Figure S2.**
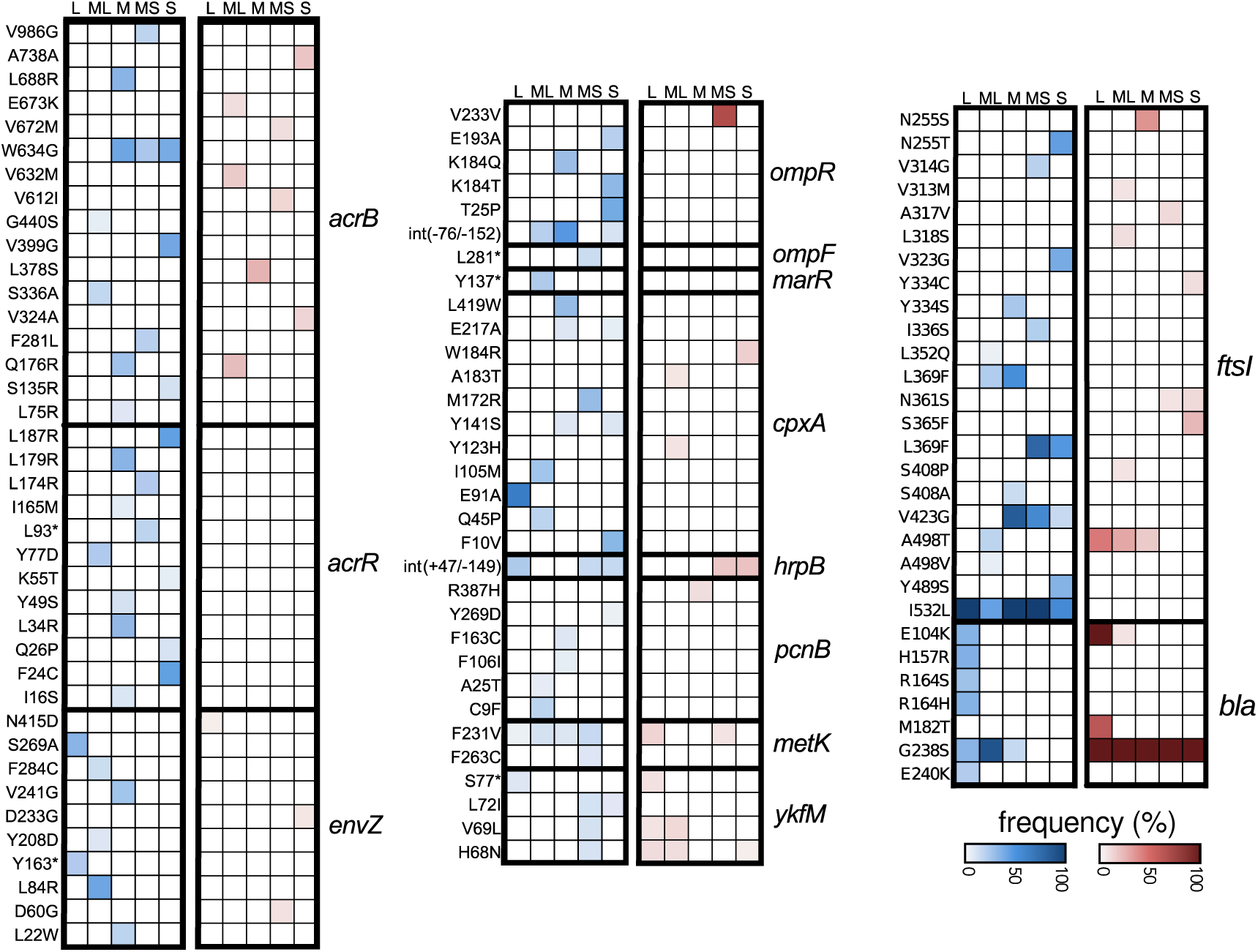
Alleles observed in focal and secondary loci from whole-genome sequencing. Heatmap shows the frequency of non-synonymous point mutations across MutT (blue) and MutH (red) populations in known genes associated with β-lactam resistance. We pooled five clones from five distinct replicates per population size treatment (columns). Darker shading indicates higher frequency of a given allele (rows) within each pool. Only 9 out of 100 mutations are shared between mutator backgrounds (6 out of 71 when excluding focal loci, *ftsI* and *bla*); all others are genotype-specific. Mutations in *acrR*, *ompF*, and *marR* occur exclusively in *MutT* populations, consistent with the need to compensate for adaptation via the “slower” PBP3 route.

**Figure S3.**
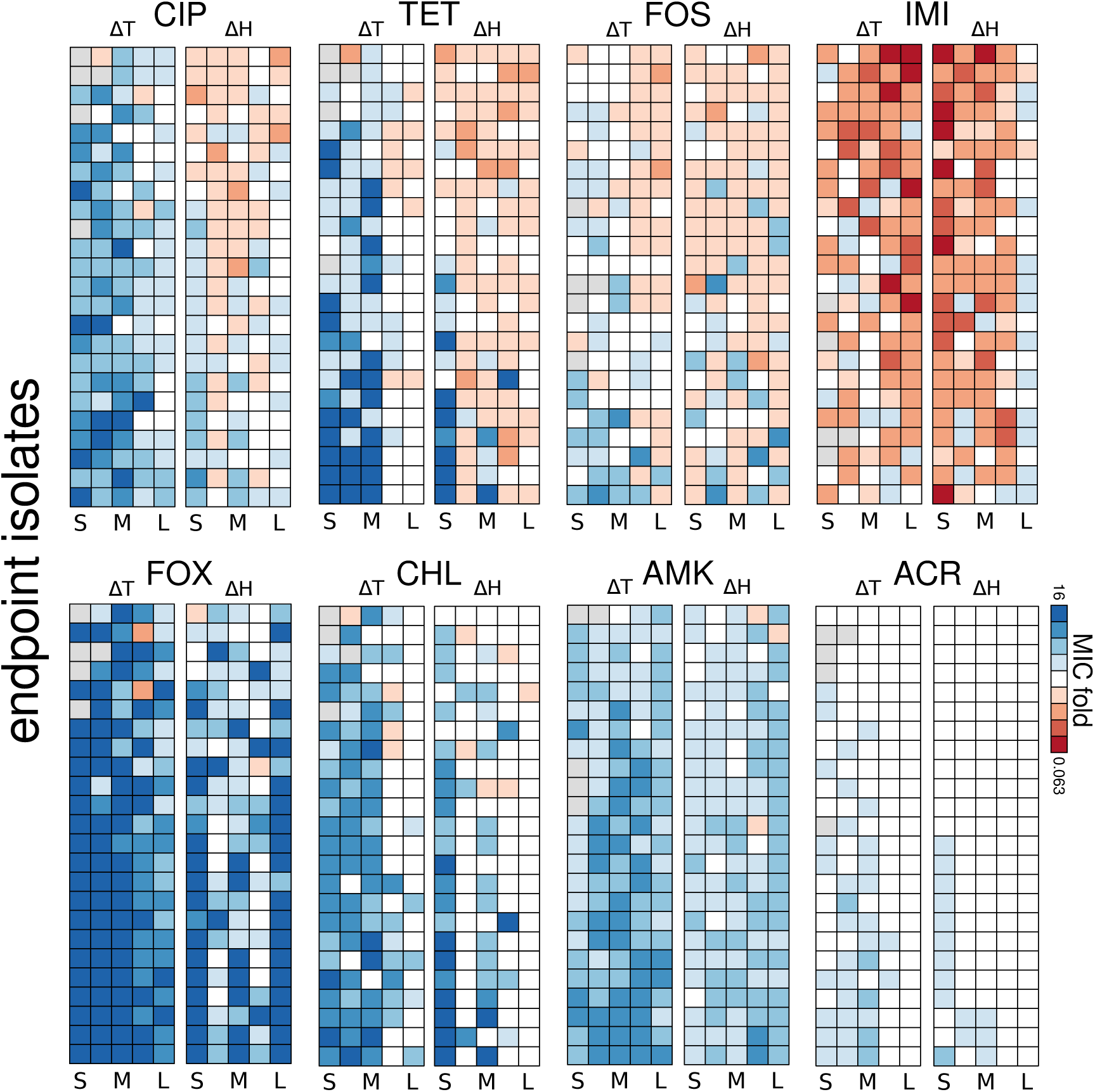
Mutators exhibit distinct collateral sensitivity patterns across population sizes. We measured collateral resistance and sensitivity to eight compounds across all treatments. Heatmaps show changes in drug susceptibility across population size classes for 24 endpoint isolates per mutator (MutT, left; MutH, right). Colors indicate fold-changes in MIC relative to the ancestor (white: no change; red: increased susceptibility; blue: increased resistance). Note that across compounds, MutT populations generally evolved higher resistance levels than MutH populations.

